# Cultivation and genomics of the first freshwater SAR11 (LD12) isolate

**DOI:** 10.1101/093567

**Authors:** Michael W. Henson, V. Celeste Lanclos, Brant C. Faircloth, J. Cameron Thrash

## Abstract

Evolutionary transitions between fresh and salt water happen infrequently among bacterioplankton. Within the ubiquitous and highly abundant heterotrophic Alphaproteobacteria order *Pelagibacterales* (SAR11), most members live in marine habitats, but the LD12 subclade has evolved as a unique freshwater lineage. LD12 cells occur as some of the most dominant freshwater bacterioplankton, yet this group has remained elusive to cultivation, hampering a more thorough understanding of its biology. Here, we report the first successful isolation of an LD12 representative, strain LSUCC0530, using high throughput dilution to extinction cultivation methods, and its complete genome sequence. Growth experiments corroborate ecological data suggesting active populations of LD12 in brackish water up to salinities of ~5. LSUCC0530 has the smallest closed genome thus far reported for a SAR11 strain (1.16 Mbp). The genome affirms many previous metabolic predictions from cultivation-independent analyses, like a complete Embden-Meyerhof-Parnas glycolysis pathway, but also provides novel insights, such as the first isocitrate dehydrogenase in LD12, a likely homologous recombination of malate synthase from outside of the SAR11 clade, and analogous substitutions of ion transporters with others that occur throughout the rest of the SAR11 clade. Growth data support metagenomic recruitment results suggesting temperature-based ecotype diversification within LD12. Key gene losses for osmolyte uptake provide a succinct hypothesis for the evolutionary transition of LD12 from salt to fresh water. For strain LSUCC0530, we propose the provisional nomenclature *Candidatus* Fonsibacter ubiquis.

## Introduction

Bacterioplankton in the SAR11 clade of *Alphaproteobacteria* are dominant heterotrophs in marine and freshwater systems. In the oceans, SAR11 can represent 25-50% of total planktonic cells (Morris *et al.*, 2002, Schattenhofer *et al.*, 2009). Numerous subclades with unique spatio-temporal distributions comprise SAR11 (Carlson *et al.*, 2009, Giovannoni and Vergin, 2012, Morris *et al.*, 2002, Thrash *et al.*, 2014). At least nine subclades defined via 16S rRNA gene sequences occupy marine niches (Giovannoni and Vergin, 2012), and more likely exist (Tsementzi *et al.*, 2016). However, in spite of its global distribution (Morris *et al.*, 2002), massive predicted population size of 10^28^ cells (Morris *et al.*, 2002), and an estimated divergence time from its last common ancestor of 1.1 billion years ago (Luo *et al.*), the bulk of existing evidence suggests that SAR11 has only successfully colonized freshwater environments once in its natural history (Eiler *et al.*, 2014, Logares *et al.*, 2010, Salcher *et al.*, 2011). Traditionally, all known freshwater SAR11 belong to subclade IIIb, a.k.a. LD12. However, a recent report challenges this assertion: a genome sister to subclade I was recovered in Lake Baikal metagenomic data (Cabello-Yeves *et al.*, 2018). Regardless, the limited evolutionary diversification into less saline habitats has not prevented LD12 from achieving prominence in the ecosystems it inhabits. In many lotic and lentic environments, LD12 occupies similar relative abundances as its marine cousins (Dupont *et al.*, 2014, Garcia *et al.*, 2017, Henson *et al.*, 2016, Newton *et al.*, 2011, Salcher *et al.*, 2011). Study of LD12 is important to understand SAR11 evolution, specifically, and how successful transitions between marine and freshwater environments occur in bacterioplankton (Logares *et al.*, 2009), more generally.

Ecological, functional, and sequence-based inference from single amplified genomes (SAGs) and metagenomes support the hypothesis that LD12 bacterioplankton evolved from a genome-streamlined marine ancestor (Eiler *et al.*, 2016, Luo *et al.*, 2013, Salcher *et al.*, 2011, Zaremba-Niedzwiedzka *et al.*, 2013). As such, they share many of the same characteristics as marine SAR11, such as small cell volumes; adaptation to oligotrophic habitats; small, streamlined genomes; an obligate aerobic chemoorganoheterotrophic lifestyle with limited metabolic flexibility; preference for small molecular weight compounds like carboxylic and amino acids as carbon/energy sources; and auxotrophies for some amino acids and vitamins (Dupont *et al.*, 2014, Eiler *et al.*, 2014, Eiler *et al.*, 2016, Giovannoni *et al.*, 2005a, Giovannoni *et al.*, 2005b, Grote *et al.*, 2012, Salcher *et al.*, 2011, Schwalbach *et al.*, 2010, Sun *et al.*, 2011, Thrash *et al.*, 2014, Tripp *et al.*, 2008, Zaremba-Niedzwiedzka *et al.*, 2013). Previous research suggests that LD12 differ from their marine counterparts in specific elements of metabolic potential that indicate a greater emphasis on production, rather than uptake, of osmolytes, and important metabolic changes related to energy production (Dupont *et al.*, 2014, Eiler *et al.*, 2016). For example, metagenomic population data showed a correlation between decreased salinity and greater proportion of the Embden-Meyerhof-Parnass (EMP) vs. Entner-Doudoroff (ED) glycolysis pathways (Dupont *et al.*, 2014). Comparative genomic analyses of SAGs from different SAR11 strains concurred: LD12 genomes contained the EMP pathway that is not found in most marine SAR11 (Eiler *et al.*, 2016, Grote *et al.*, 2012). SAG data also suggested that LD12 lacks the glyoxylate shunt and some single carbon (C1) metabolism (Eiler *et al.*, 2016).

Despite what has been learned from cultivation-independent methods, the lack of cultured LD12 representatives has hampered a more detailed exploration of the group. Potential ecotypes within LD12 have been identified (Zaremba-Niedzwiedzka *et al.*, 2013), and their population dynamics recently described with five-year time series data in freshwater lakes (Garcia *et al.*, 2017). However, we cannot delineate what distinguishes ecotypes without better physiological and genomic data. Similarly, interpreting data on the ecological distribution of LD12 remains challenging without information on growth tolerances and optima for salinity and temperature. We also don’t understand whether a connection exists between more efficient energy production through EMP-based glycolysis and the freshwater lifestyle, or what other adaptations might explain LD12 evolution away from salt water.

The next steps in translating ‘omics-based predictions into measured data for integration with ecosystem models require living experimental subjects. For example, cultures of marine SAR11, such as HTCC1062 and HTCC7211, have facilitated testing of metabolism and growth (Carini *et al.*, Carini *et al.*, Giovannoni *et al.*, 2005a, Schwalbach *et al.*, 2010, Smith *et al.*, Smith *et al.*, Steindler *et al.*, 2011, Tripp *et al.*, 2008), structural organization (Zhao *et al.*, 2017), and virus-host interactions (Zhao *et al.*, 2013). We need cultivated representatives to provide this kind of understanding of other important bacterioplankton like LD12. In service of this goal, we pursued a systematic high throughput cultivation effort from coastal regions in the northern Gulf of Mexico. Here, we report the first isolation of a member of the LD12 clade, strain LSUCC0530, its complete genome sequence, preliminary physiological data, and an examination of the ecological distribution of this organism compared with other LD12 clade members. Our results provide evidence for temperature-dependent ecotype diversification within LD12 and a hypothesis about the evolutionary trajectory that led to LD12 colonization of fresh water.

## Results

### Isolation and growth of LSUCC0530

We isolated strain LSUCC0530 as part of a series of high throughput dilution-to-extinction cultivation experiments (HTC) conducted with inoculation water obtained across the southern Louisiana coast. The particular experiment that yielded LSUCC0530 utilized surface water from the coastal lagoon of Lake Borgne, 39 km southeast of New Orleans. At the time of collection, the surface water had a salinity of 2.39 and was 30.5°C. Subsequent community analysis demonstrated that LD12-type organisms represented 8.7% of the bacterioplankton population in the inoculum (see below). We diluted 2.7 μm-filtered Lake Borgne surface water for inoculation to an estimated 2 cells well^-1^ into our JW5 medium (salinity 1.45, Table S1), and incubated cultures for 30 days at 23°C (room temperature in the Thrash Lab). LSUCC0530 grew slowly, reaching 8.6 x 10^5^ cells mL^-1^ after 24 days. It had a low fluorescence/low side scatter flow cytometric signature characteristic of other SAR11 cells (Fig. S1).

### Taxonomy and morphology

The LSUCC0530 16S rRNA gene sequence had > 99% identity with eight SAG sequences from members of LD12 (Zaremba-Niedzwiedzka *et al.*, 2013) and only 91% and 92% sequence identity with sequences from the sister subclade IIIa representatives IMCC9063 and HIMB114, respectively. Of note for taxonomic incorporation of genomic data (Chun *et al.*, 2018), the genome-generated 16S rRNA gene sequence matched the PCR-generated 16S rRNA gene sequence (GenBank accession number KY290650.1) at 1316 of 1320 sites. Two of these mismatches were single bp gaps the genome sequence, and three of the four mismatches occurred in the first 20 bp of the PCR-generated sequence. LSUCC0530 also had average amino acid identities (AAI) of 58.1% and 59.7% with HIMB114 and IMCC9063, respectively (Table S1). In contrast, AAI between LSUCC0530 and the LD12 SAGs ranged from 81.9% to 88.0%, corroborating that LSUCC0530 belongs to separate species from subclade IIIa (Konstantinidis and Tiedje, 2007). Phylogenetic inference using 16S rRNA genes placed LSUCC0530 within the Order *Pelagibacteriales* (a.k.a. SAR11 (Thrash *et al.*, 2014)), in monophyly with the original LD12 clone library sequence (Fig. S2). Phylogenomic inference using 83 single copy marker genes from 49 SAR11 genomes also strongly supported placement of LSUCC0530 within LD12 as an early diverging member, with only the SAG AAA280-B11 from Lake Damariscotta diverging earlier (Fig. 1A). This tree also recovered the three microclusters within LD12 (A, B, C) observed previously (Zaremba-Niedzwiedzka *et al.*, 2013) (Fig. 1A). Cells of strain LSUCC0530 were small, curved rods, approximately 1 μm x 0.1 μm (Fig. 1B, C), showing conserved morphology with the distantly related SAR11 strain HTCC1062 in subclade Ia (Rappé *et al.*, 2002, Zhao *et al.*, 2017).

**Figure 1.**
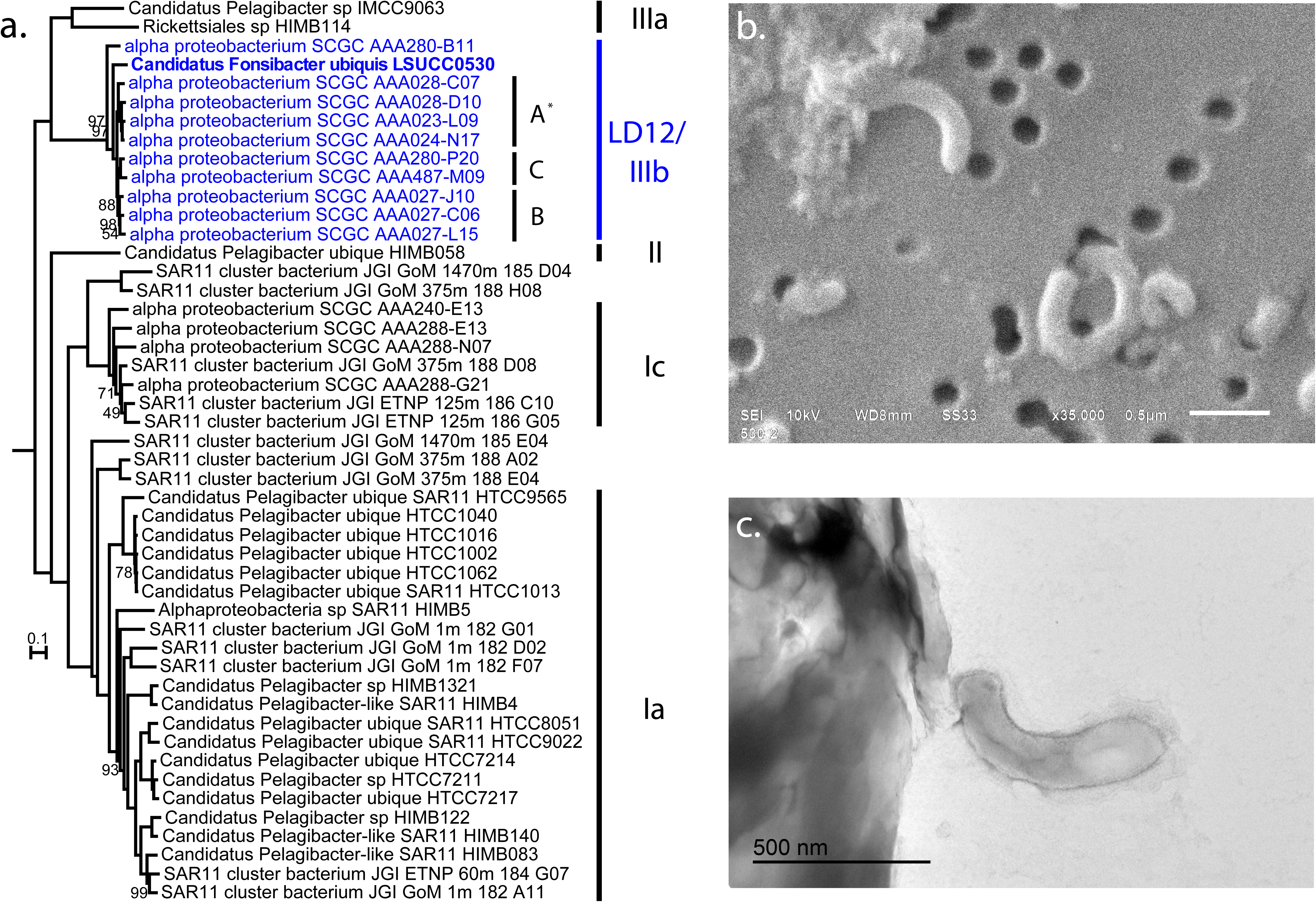
Phylogeny and morphology of LSUCC0530. **A)** Phylogenetic position of LSUCC0530 within the SAR11 clade using a concatenation of 83 single copy marker genes and 23,356 amino acid positions. The tree was inferred using RAxML with 100 bootstrap runs. Values at the nodes indicate 100% bootstrap support unless otherwise noted. The tree was rooted on HIMB59. Subclades are indicated on the right, and LD12 microclusters are indicated A-C. Asterisk - we have included AAA028-C07 in microcluster A, even though it was excluded previously (Zaremba-Niedzwiedzka *et al.*, 2013). **B)** Scanning electron micrograph of LSUCC0530 cells at 35,000x magnification. Scale bar represents 0.5 μm. **C)** Transmission electron micrograph of a LSUCC0530 cell thin section. Scale bar represents 0.5 μm.

### Genome characteristics

Genome assembly resulted in a single, circularized scaffold that we tested for completeness using multiple tools (Supplemental Information). The completed genome is 1,160,202 bp with a GC content of 29.02% and 1271 predicted genes (Fig. 2). There are 1231 putative protein coding genes and 40 RNA genes, including one each of the 5S, 16S, and 23S rRNA genes. We found no genomic evidence of lysogenic bacteriophage or a CRISPR-*cas* system. LSUCC0530 has the characteristics of genome streamlining previously reported for other SAR11 genomes, with an estimated coding percentage of 96.36% and the smallest complete SAR11 genome reported thus far (Giovannoni *et al.*, 2005b, Grote *et al.*, 2012). Despite its smaller overall genome size, intergenic spacer regions are not significantly smaller than those of the sister subclade IIIa (Fig. S3). All LD12 genomes had much higher AAI and synteny with each other than with the subclade IIIa (Table S1), mirroring their phylogenomic relationships (Fig. 1A). However, within the LD12 clade, AAI and synteny percentages did not clearly delineate the same patterns we observed in the phylogenomic tree, which may have resulted from the LD12 SAGs being fragmented and incomplete.

**Figure 2.**
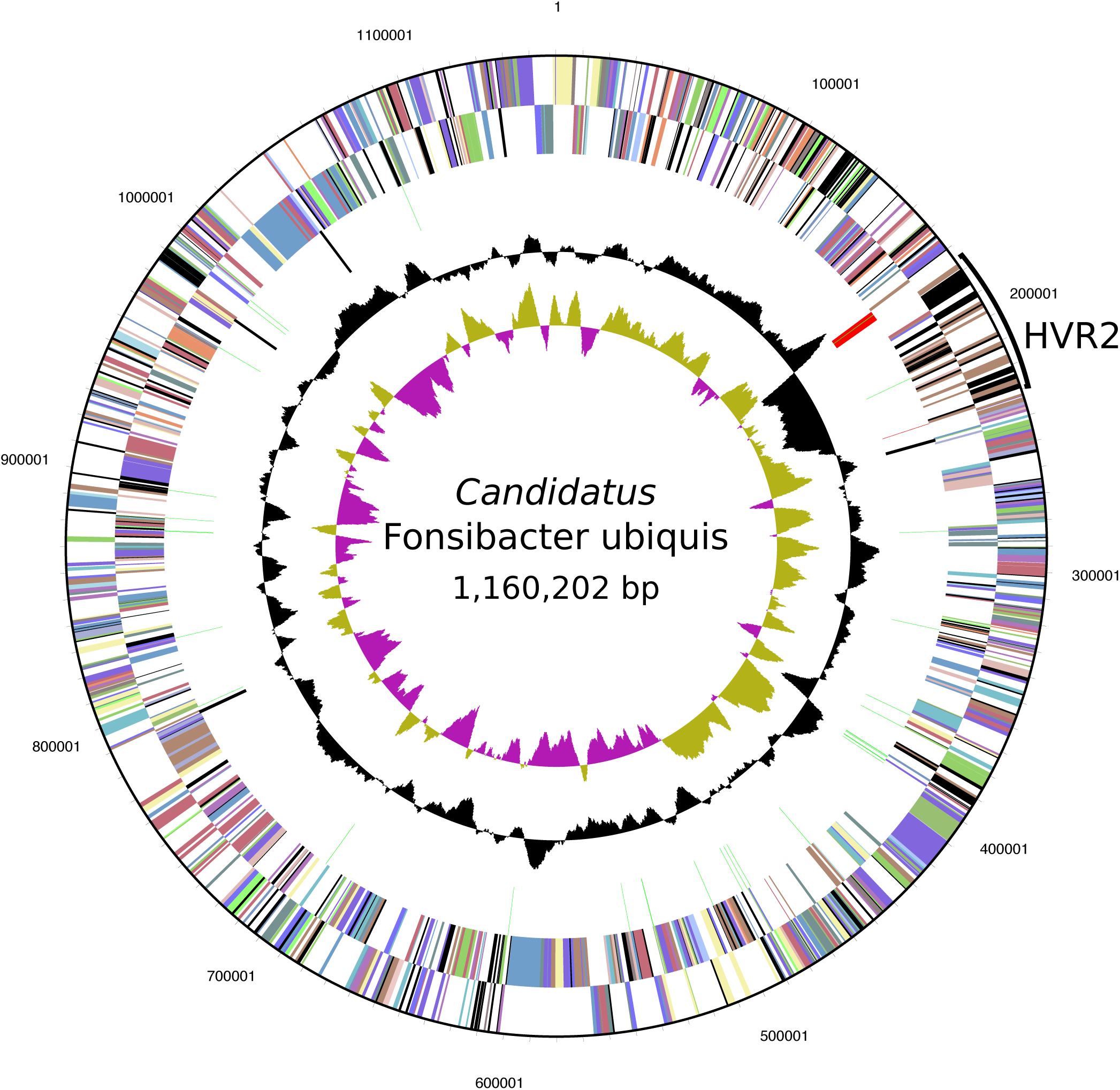
Circular diagram of the LSUCC0530 genome. Rings indicate, from outside to the center: Position relative to the replication start site; Forward strand genes, colored by COG category; Reverse strand genes, colored by COG category; RNA genes (tRNAs green, rRNAs red, other RNAs black); GC content; GC skew.

The LSUCC0530 genome has the conserved architecture for hypervariable region 2 (HVR2) that has been observed in most other complete SAR11 genomes (Grote *et al.*, 2012, Rodriguez-Valera *et al.*, 2009, Wilhelm *et al.*, 2007) and other LD12 SAGs (Zaremba-Niedzwiedzka *et al.*, 2013), between the 16S rRNA-tRNA-tRNA-23S rRNA operon and the 5S rRNA gene operon. This region is 54,755 bp and has a strong variation in GC content from neighboring parts of the genome (Fig. 2). Metagenomic recruitment also verified this region as hypervariable based on the lack of matching sequences in community data (Fig. S4). The predicted gene annotations in HVR2 resemble those in the same location in other SAR11 genomes (Grote *et al.*, 2012, Zaremba-Niedzwiedzka *et al.*, 2013), namely genes likely functioning in membrane modification such as glycosyltransferases, epimerases, transaminases, and methyltransferases; as well as hypothetical proteins.

### Metabolic reconstruction

Like most other SAR11s (Eiler *et al.*, 2016, Grote *et al.*, 2012, Thrash *et al.*, 2014), the LSUCC0530 genome encodes for an obligate aerobic lifestyle, with a complete oxidative phosphorylation pathway including a heme/copper-type cytochrome c oxidase, a proton-translocating NADH dehydrogenase, and a proton-translocating ATP synthase (Fig. 3). As previously detected in LD12 SAGs (Eiler *et al.*, 2016), the LSUCC0530 genome encodes a complete EMP glycolysis pathway, including phosphofructokinase and pyruvate dehydrogenase, and complete gluconeogenesis through nucleotide sugar production (Fig. 3). We recovered genes for the pentose phosphate pathway and a predicted fructokinase gene for conversion of D-fructose to ß-D-fructose-6-P. As expected based on other SAR11 genomes, we did not find annotated genes for utilization of allose, fucose, fuculose, galactose, maltose, mannose, rhamnose, sucrose, starch, trehalose, or xylose. LSUCC0530 has a complete TCA cycle and glycolate oxidase, and like other SAR11s, the non-mevalonate pathway for isopentenyl diphosphate (Fig. 3).

**Figure 3.**
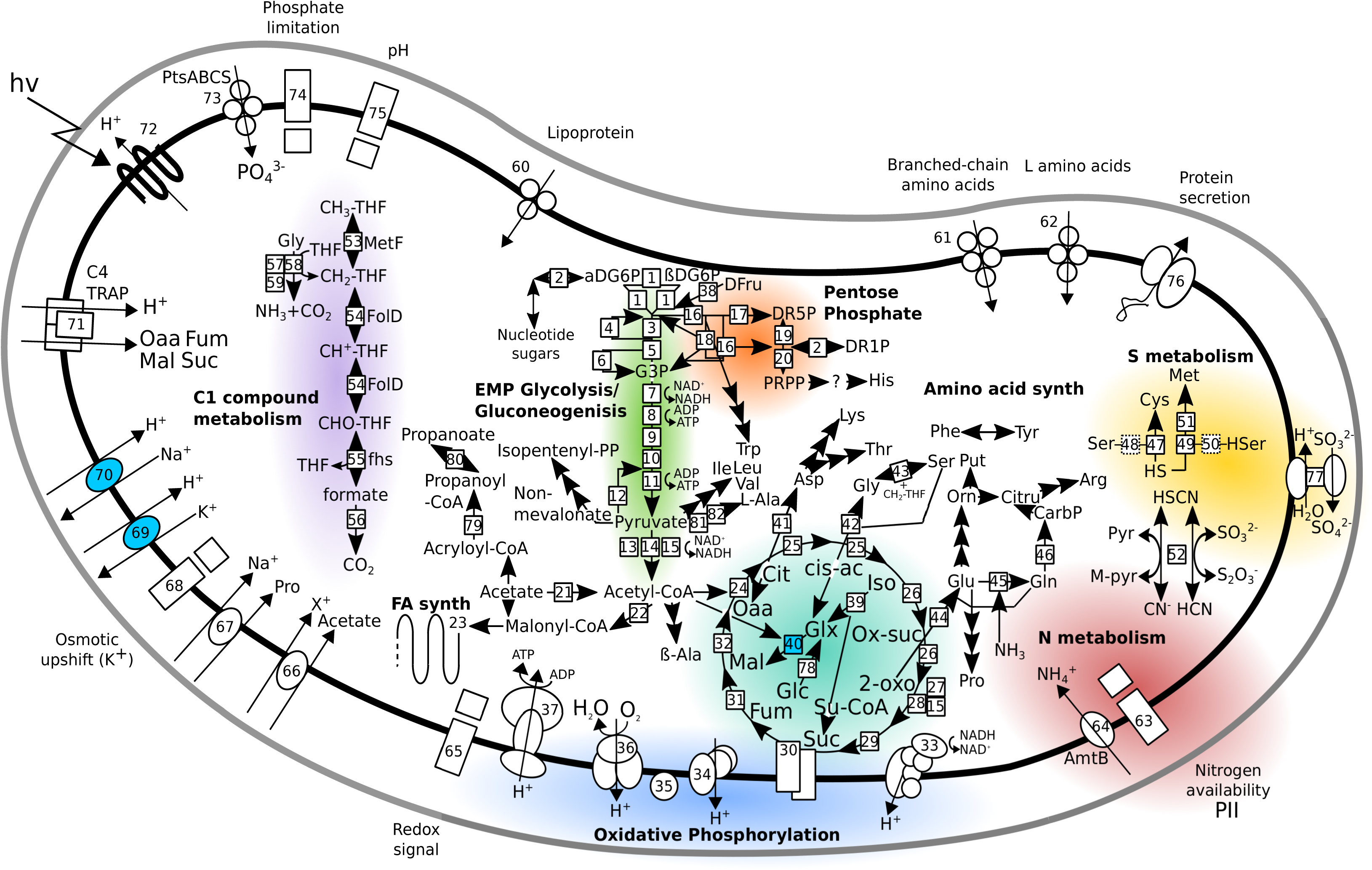
Metabolic reconstruction of LSUCC0530. Solid boxes indicate recovered genes, dashed boxes indicate missing genes. Color shading distinguishes major metabolic subsystems, which are also labeled in bold text. ABC transporters are depicted with circles indicating subunits. Symporters and antiporters are depicted with ovals. Two component systems are depicted with large and small rectangles. The C4 TRAP transporter is depicted separately. Major subsystems are colored for easier identification. Multiple arrows indicate all genes present in a given pathway. Numbers identify genes according to the key in Table S1. Question marks indicate a single missing gene in an otherwise complete pathway (e.g., PRPP -> His). Light blue fill indicates genes with no SAR11 orthologs outside of the LD12 subclade.

We recovered the first isocitrate lyase in an LD12 genome, conferring a complete glyoxylate bypass in combination with malate synthase. The isocitrate lyase shares its ancestry with the majority of the SAR11 clade except subclade IIIa, where it appears to have been lost (Eiler *et al.*, 2016, Grote *et al.*, 2012) (Fig. S5). However, the LSUCC0530 malate synthase shared with other LD12 SAGs belongs to isoform A, whereas all other SAR11 clades contain isoform G (Anstrom *et al.*, 2003) (Figs. 3; S6). Comparison of gene neighborhoods in a variety of SAR11 genomes revealed that both isoforms occur in the same location (Fig. S7).

We predict production of eighteen amino acids (Fig. 3) and partial synthesis pathways for three more (histidine, cysteine, and methionine), as well as the ability to interconvert, but not synthesize *de novo*, phenylalanine and tyrosine. The LSUCC0530 genome has lost some of the C1 and methylated compound pathways found in other SAR11s (Sun *et al.*, 2011). For example, as in the LD12 SAGs (Eiler *et al.*, 2016), LSUCC0530 does not have DMSP, methylamine, or glycine-betaine metabolism. However, it retains tetrahydrafolate metabolism, formate oxidation, and the glycine cleavage pathway (Fig. 3).

The genome has the same PII nitrogen response system and *amtB* ammonium transporter found in other SAR11s and is predicted to utilize ammonia as a nitrogen source in amino acid synthesis (Fig. 3). The genome also encodes a putative thiosulfate/3-mercaptopyruvate sulfurtransferase that may be involved in sulfur and thiocyanate reactions (Fig. 3). A possible product of this enzyme is sulfite, which could be further oxidized to sulfate in the periplasm via putative *sorAB* genes. This gene pair occurs infrequently in the SAR11 clade, but sulfite oxidation may serve, in some strains, as a means to generate additional proton motive force (Denger *et al.*, 2008, Kappler, 2011). This is similar to the function conferred by proteorhodopsin (Béjà *et al.*, 2001, Giovannoni *et al.*, 2005a, Steindler *et al.*, 2011), which the LSUCC0530 genome also contains. Also like other SAR11s, a complete assimilatory sulfate reduction pathway is missing, making LSUCC0530 dependent on reduced sulfur compounds for cellular sulfur requirements (Tripp *et al.*, 2008). In our media, this was supplied in the form of sulfur-containing amino acids (Table S1). The genome encodes partial pathways for incorporation of sulfide into cysteine and methionine but is missing predicted serine O-acetyltransferase and homoserine O-succinyltransferase genes (Fig. 3).

In addition to the PII nitrogen sensor, LSUCC0530 also has putative two component systems handling phosphate limitation, pH, osmotic upshift via potassium, and redox state (Fig. 3). The LSUCC genome encodes ABC transporters for phosphate (*ptsABCS*), lipoprotein, and branched-chain and L amino acid uptake. We identified tripartite ATP-independent periplasmic (TRAP) transporter genes for C4-dicarboxylate and mannitol/chloroaromatic compounds. However, the gene cluster appears to be missing the substrate binding subunit, and we did not find predicted genes for mannitol utilization.

In general, LSUCC0530 and the LD12 SAGs share many of the same pathways for osmolyte synthesis as the subclade IIIa and other SAR11 organisms, such as N-acetyl-ornithine, glutamate, glutamine, and proline (Table 1). In some cases, there appear to be analogous substitutions in osmolyte transport systems. For example, LD12 genomes have potassium antiporters, but not the *trkAH* genes for potassium transport in other SAR11s. Similarly, a sodium/proton antiporter orthologus cluster comprised genes exclusively from LD12 genomes, but a separate sodium/proton antiporter occurred in all other SAR11 genomes (Table 1). Perhaps the most important difference in genomic content concerning osmolytes occurs in two key transporter losses: LSUCC0530 and the other LD12s have no genes for the glycine betaine/proline or ectoine/hydroxyectoine ABC transporters that are present in HIMB114 and IMCC9063 and other subclades of SAR11 (Table 1).

**Table 1.** Compatible solute production and transport in SAR11

| Osmolyte system | LSUCC0530 | LD12 SAGs | HIMB114 | IMCC9063 | Other SAR11 |
| --- | --- | --- | --- | --- | --- |
| Alanine synthesis | + | + | + | + | + |
| Di-myo-inositol phosphate synthesis | +/- | +/- | +/- | +/- | +/- |
| Ectoine/hydroxyectoine transport | - | - | + | + | + |
| Glutamate synthesis | + | + | + | + | + |
| Glutamate transport | - | + | - | - | - |
| Glutamine synthesis | + | + | + | + | + |
| Glycine synthesis | + | + | + | + | + |
| Glycine betaine synthesis | - | +/-* | + | + | + |
| Glycine betaine/proline ABC transporter | - | - | + | + | + |
| N-δ-Acetyl-ornithine synthesis | + | + | + | + | + |
| Potassium transport (antiporters K03549, K03455) | + | + | - | - | - |
| Potassium transport (trkAH + K16052) | - | - | + | + | + |
| Proline synthesis | + | + | + | + | + |
| Proline transport (symporter) | + | + | + | + | + |
| Sodium transport (antiporter K03313) | - | - | + | + | + |
| Sodium transport (other antiporter) | + | + | - | - | - |
| Sorbitol synthesis | - | - | + | - | + |
| Taurine transport | - | - | - | - | + |
| TMAO synthesis | - | - | - | - | + |

### Physiology

Under isolation conditions (23°C, JW5 medium), strain LSUCC0530 grew to a density of 2 x 10^7^ cells mL^-1^ with an average growth rate of 0.52 day^-1^. Our physiological experiments to determine the salinity tolerance and growth optima revealed that LSUCC0530 could grow at salinities between 0.36 and 4.7 (Fig. 4A). The optimal salinity for LSUCC0530 was at or below the isolation medium of 1.45. Growth rates decreased between salinities of 2.9 and 4.7 (Fig. 4A), and cells died (determined by rapid loss of flow cytometric signal) in salinities above 8 (Fig. S8). We observed growth after a very extended lag phase in a subset of cultures at salinity 5.8, but could not always reproduce this growth in repeated experiments (Fig. S8). Among the temperatures tested, LSUCC0530 grew optimally at 23°C, and we observed growth up to 30°C, but not at 12°C or below, nor at 35°C or above (Fig. 4B). However, we did not observe a clear loss of signal at 4, 12, 35, or even 40°C (Fig. S8), even after over 33 days, suggesting that LSUCC0530 can endure extended periods at these temperatures in a non-growth state. Notably during growth experiments, one of the five replicates during any given experiment frequently grew at a reduced rate than the others (Fig. S8).

**Figure 4.**
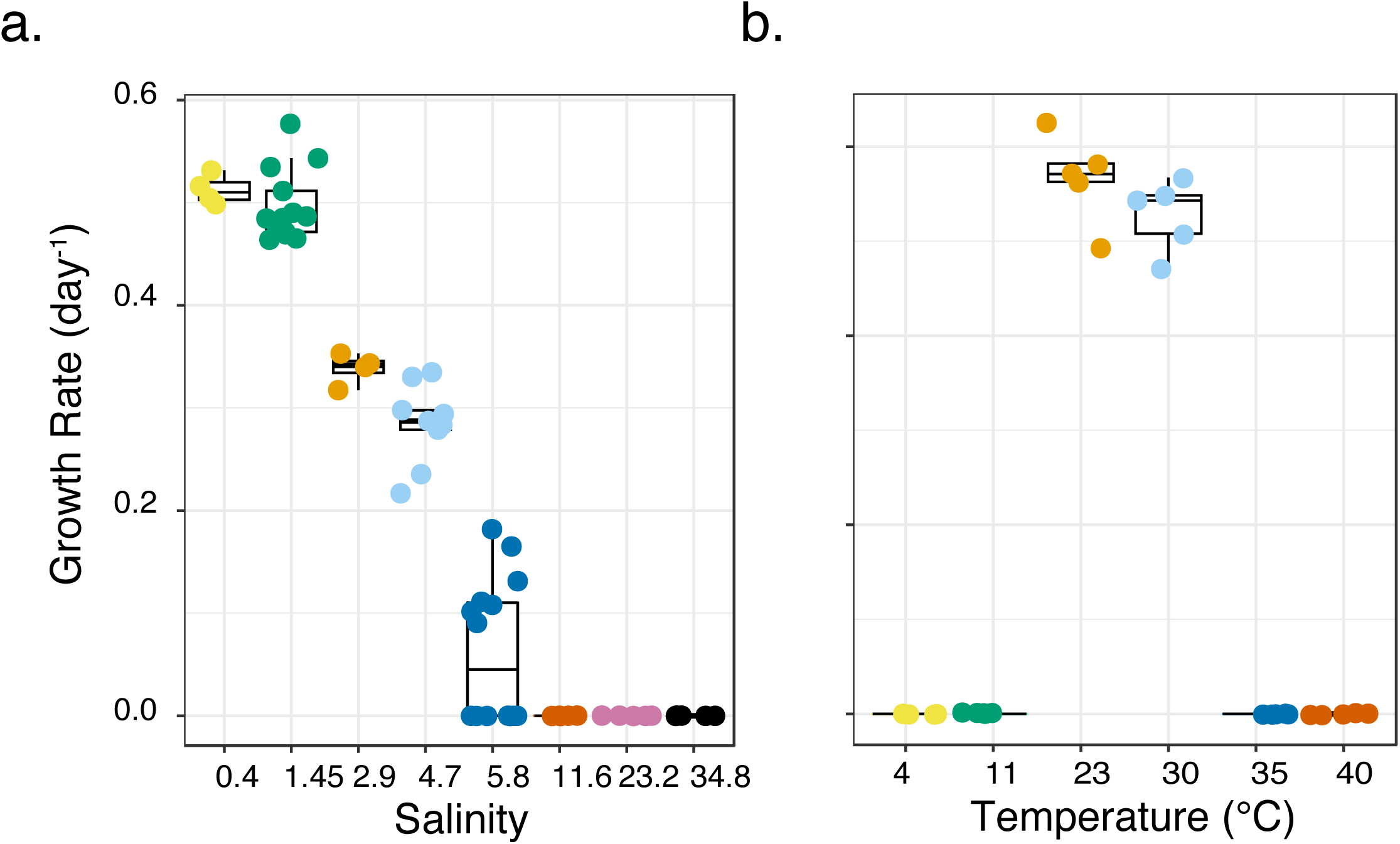
Growth rates for LSUCC0530 according to **A)** salinity and **B)** temperature. The boxes indicate the interquartile range (IQR) of the data, with vertical lines indicating the upper and lower extremes according to 1.5 x IQR. Horizontal lines within each box indicate the median. The underlying data points of individual biological replicates, calculated from corresponding growth curves in Figure S8, are plotted on top of each box.

### Ecology

We quantified the abundance of LD12 taxa in the coastal Louisiana ecosystems using 16S rRNA gene sequences clustered into operational taxonomic units (OTUs). These sequences were obtained as part of the combined sampling and HTC campaign that included the successful isolation of LSUCC0530. Based on BLASTn sequence similarity, the representative sequence for OTU7 and LSUCC0530 shared 100% identity over the 250bp v4 region of the 16S rRNA gene, confirming the Mothur classification of OTU7 as belonging to the SAR11 LD12 clade. Over the three-year sampling period, OTU7 was the fifth most abundant OTU based on total read abundance and sixth ranked OTU based on median read abundance. At the time of sampling for the experiment that resulted in the isolation of LSUCC0530, OTU7 had a relative abundance of 8.7% in Lake Borgne. In some sites, such as the Atchafalaya River Delta (salinity 0.18), LD12 represented as much as 16% of total bacterioplankton (Fig. S9). Generally, the LD12 OTU7 relative abundance correlated negatively with salinity. At seven sites with salinities below 6, OTU7 had a relative abundance of 4.0% or greater with a median relative abundance of 8.8% (Fig. S9). At the remaining ten sites above a salinity of 6 (median 19.44), OTU7 had relative abundances around 2% or lower (Fig. S9).

The phylogenetic separation of LSUCC0530 from B11 and the IIIb A-C microclusters raised the question of whether LSUCC0530 represents a novel ecotype with unique spatiotemporal distribution compared to other LD12 taxa. Therefore, we explored the distribution of LD12 genomes in 90 fresh and brackish water environments through competitive recruitment of metagenomic sequences to all the SAR11 genomes in the phylogenomic tree and the outgroup sequence HIMB59. We quantified relative abundance via recruited reads (per kpb of metagenomic sequence per Mbp of genome sequence-RPKM), and aggregated LD12 data based on the A, B, and C microclusters (Fig. 1A), the B11 SAG, and LSUCC0530. Figure 5A displays RPKM values for reads with ≥ 95% identity, roughly corresponding to a species delineation via average nucleotide identity (ANI)(Konstantinidis and Tiedje, 2005), but we also provide additional comparisons at ≥ 85, 90, 92, 98, and 100% identity (Fig. S10).

**Figure 5.**
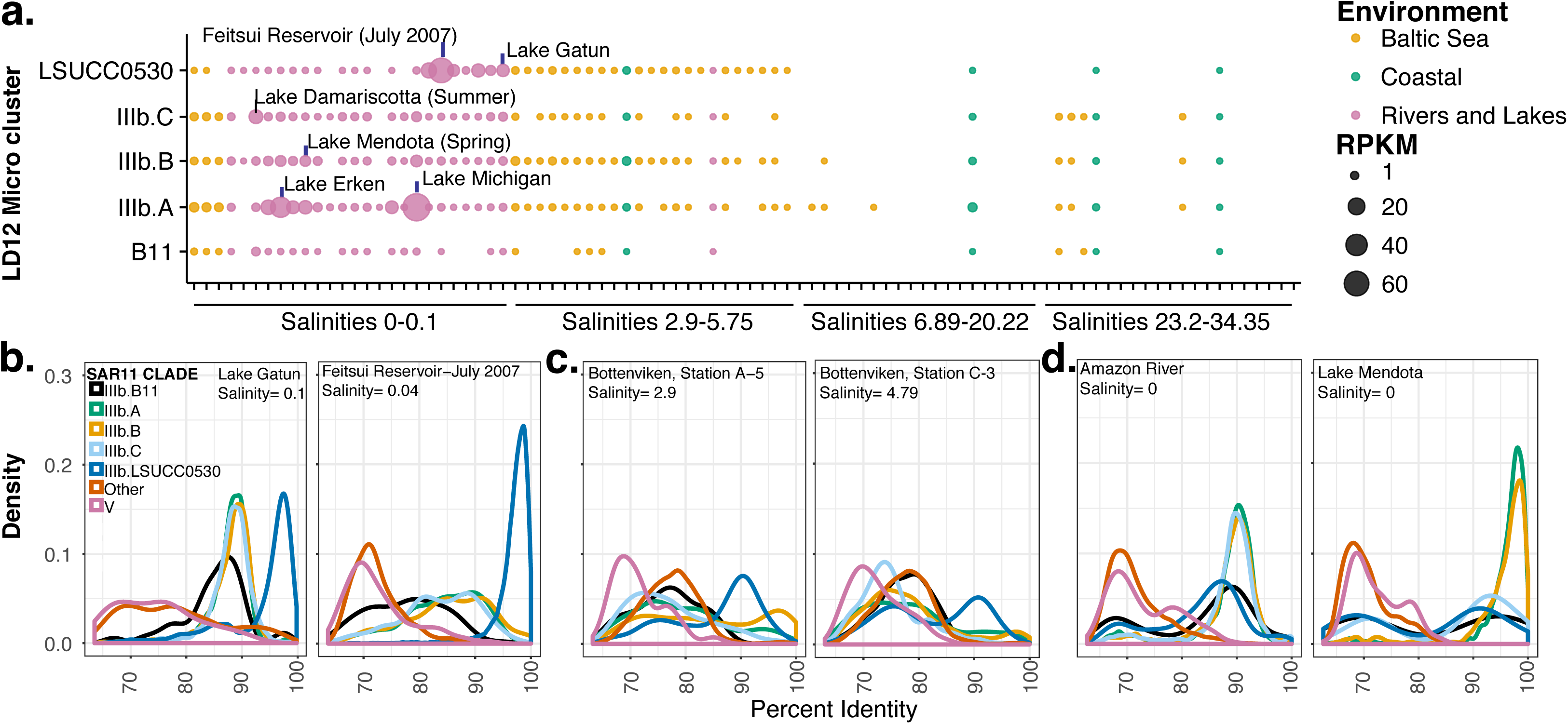
Results of competitive metagenomic recruitment to all genomes used in the phylogenomic tree. **A)** 95% identity RPKM values for LD12 genomes only, by site, with data aggregated for all genomes in microclusters A-C, according to the key. Colors indicate broad environmental categories. **B-D)** Density plots of metagenomic sequence recruitment according to percent identity of the best hit. Data was aggregated according to the key, with LD12 separated as in A, HIMB59 separately depicted as “V”, and all other SAR11 genomes together as “Other.” **B)** Two sites where LSUCC0530 dominated recruitment with high percent identity. **C)** Two sites where the LSUCC0530 only recruited sequences at ~90% identity and no other group had high recruitment at higher identity. **D)** Two sites where other LD12 microclusters showed similar patterns as LSUCC0530 in B and C.

All LD12 subgroups had their highest RPKM values in very low salinity sites, although we could detect LD12 across various brackish salinities (Fig. 5A). In general, IIIb.A and LSUCC0530 averaged the highest relative abundances across all sites with salinities between 0 and 5.75. All subgroups showed site-specific variation in abundances (Fig. 5A). Microcluster IIIb.A clearly dominated recruitment at Lake Michigan, whereas LSUCC0530 represented the most abundant genome at Feitsui Reservoir and Lake Gatun (Fig. 5A).

We repeated the previous examination of recruitment density to the different microclusters across a range of percent identities, now including LSUCC0530 (Garcia *et al.*, 2017). We corroborate observed patterns indicating additional LD12 diversity beyond that represented in the current genomes, despite the inclusion of our new representative. For example, whereas the LSUCC0530 genome recruited the majority of high percent identity reads in the Feitsui Reservoir and Lake Gatun samples (Fig. 5B), in some locations the majority of recruitment to this genome centered around lower percent identity hits (Fig. 5C). We observed the same kind of alternative patterns for the other microclusters as well (Fig. 5D). As noted our colleagues, since reads were competitively recruited against all close relative genomes, these cases where a microcluster primarily recruited reads ~90-92% identity likely mean that we lack a representative genome for that fraction of the metagenomic data (Garcia *et al.*, 2017).

## Discussion

Our successful cultivation of an LD12 strain for the first time has demonstrated that this organism shares many physiological traits with cultured marine SAR11 representatives, and also has some important distinctions. LSUCC0530 doubles roughly once every two days, similar to taxa in subclade Ia (Rappé *et al.*, 2002) and Ib (Jimenez-Infante *et al.*, 2017). It has an aerobic, chemoorganoheterotrophic lifestyle, thriving in oligotrophic media with simple carbon compounds and likely depends on reduced organosulfur compounds and ammonium as its sulfur and nitrogen sources, respectively. Cell size and shape showed considerable morphological conservation (small, curved rods roughly 1 x 0.1 μm) across large evolutionary distances (LD12 vs. subclade Ia ~89% 16S rRNA gene identity). Although a systematic survey of salinity tolerance has not been conducted for other cultured SAR11 representatives, we have shown that LD12 cannot survive in media with salinities over eight, whereas all other cultured SAR11 grow in seawater salinities (Carini *et al.*, 2013, Jimenez-Infante *et al.*, 2017, Rappé *et al.*, 2002, Song *et al.*, 2009, Stingl *et al.*, 2007). LSUCC0530 thrives in relatively high temperatures (23-30°C), and we could not confirm growth at low temperatures in which other SAR11 have been isolated (Song *et al.*, 2009).

LSUCC0530 shares many genomic characteristics with marine SAR11, such as a small, streamlined genome with few paralogs, short intergenic spaces, and a conserved hypervariable region, confirming predictions from LD12 SAGs and metagenomes (Eiler *et al.*, 2016, Newton *et al.*, 2011, Zaremba-Niedzwiedzka *et al.*, 2013). The LSUCC0530 genome also encodes much of the same metabolic potential as what has been reported through LD12 SAGs and metagenomic data. For example, we confirm the EMP glycolysis pathway, TCA cycle, non-mevalonate terpenoid biosynthesis, numerous amino acid synthesis pathways, TRAP transporters for C4 dicarboxylate compounds, and ABC transporters for various amino acids. We can confirm the lack of a phosphoenolpyruvate carboxylase (ppc) and missing pathways for DMSP, glycine-betaine, and methylamine metabolism.

We have also recovered important genetic information previously unknown for LD12. At 1.16 Mbp, LSUCC0530 has a smaller genome, with fewer genes, than any other SAR11. LSUCC0530 has a complete glyoxylate bypass, which differs from previous inferences using SAGs (Eiler *et al.*, 2016). The unusual LD12 malate synthase isoform A gene occupies the same genomic location in LSUCC0530 as the isoform G copy in most subclade Ia, Ic, and IIIa genomes, with the exception of HIMB114 (Fig. S7). Therefore, the most parsimonious explanation for the presence of isoform A in LSUCC0530 is a homologous recombination event that replaced the isoform G homolog. The LSUCC0530 genome also contains putative pathways for oxidation of sulfide, thiosulfate, and sulfite, which have not yet been explored in any cultured SAR11.

The relative abundance data from coastal Louisiana and the isolation of an LD12 strain from a coastal environment supports previous work indicating that LD12 occupies a much wider range of aquatic habitats than inland freshwater systems (Dupont *et al.*, 2014, Herlemann *et al.*, 2014, Piwosz *et al.*, 2013, Salcher *et al.*, 2011). Questions have been raised as to whether these organisms occur in brackish waters as active and growing community members, or whether there are present simply as a result of freshwater input. LD12 cells have been shown to uptake thymidine in brackish water (Piwosz *et al.*, 2013). Our observation that LSUCC0530 can grow in slightly brackish water provides further evidence that such habitats may support LD12 cells as active community members. However, the relative competitiveness of LD12 within brackish microbial assemblages requires further research.

Metagenomic recruitment provides evidence that LD12 has diversified into multiple ecotypes with unique distributions (Fig. 5A) (Garcia *et al.*, 2017, Zaremba-Niedzwiedzka *et al.*, 2013). Furthermore, our data suggests that LSUCC0530 likely represents a novel LD12 ecotype because it preferentially recruits metagenomic sequences at high percent identity in different locations than the other microclusters of LD12 (Fig. 5). What distinguishes the distribution of these ecotypes remains unclear. The observed patterns may represent temporal variation captured in these sampling snapshots (Garcia *et al.*, 2017). Another possibility is that small variations in salinity optima for different ecotypes may drive diversification. However, given the paucity of low-brackish sites in available datasets, we cannot establish whether salinity plays a role in ecotype diversification. Furthermore, without complete genomes from other LD12 ecotypes, we cannot rule out whether variant ionic strength management strategies, or some other metabolic characteristic, help define evolutionary differences associated with ecotypes. For example, the ability to utilize different sulfur sources may distinguish LSUCC0530 from other LD12. The *sorAB* sulfite dehydrogenase and the thiosulfate/3-mercaptopyruvate sulfurtransferase genes have uneven distribution among SAR11 genomes, including within the LD12 subclade.

An alternative hypothesis is that temperature has driven diversification between ecotypes. A striking feature of LSUCC0530 is its growth preference for relatively high temperatures and inability to grow at low temperatures. While robust growth at 30°C may seem intuitive given that this represents typical Lake Borgne surface water temperature, the inability to grow at 12°C or below would likely place LSUCC0530-type LD12 cells at a fitness disadvantage in places like Lake Michigan, where annual water temperatures rarely get above 22°C and average temperatures remain below 15°C for the majority of the year (https://coastwatch.glerl.noaa.gov/). The metagenomic recruitment data support the temperature hypothesis: the samples in which LSUCC0530 dominated recruitment (Lake Gatun, Feitsui Reservoir) were all collected from locations having average temperatures above 19°C. Conversely, other microclusters dominated recruitment of samples from colder environments (e.g. IIIb.A in Lake Michigan)(Fig. 5). More sequence data may strengthen the relationship between temperature and ecotypes, but since temperature optima are extremely hard to infer from genomes alone, additional cultivars will be needed to test the temperature hypothesis for ecotype diversification.

A critical question remains for LD12. What has made these cells uniquely adapted to low salinity waters? Genomic analyses here and elsewhere (Dupont *et al.*, 2014, Eiler *et al.*, 2016) have helped define the differences in LD12 metabolic potential compared to other SAR11, but very few adaptations appear related to salinity. It has been noted that the EMP pathway provides more ATP and NADH than the ED pathway, thus conferring greater energy conservation through glycolysis for LD12 cells than other SAR11s (Dupont *et al.*, 2014, Eiler *et al.*, 2016). If or how this energy conservation relates to salinity adaptation has not been pursued. The presence of unique LD12 cation-proton antiporters (Fig. 3) could provide a connection. LD12 cells may benefit from additional NADH for the production of a proton motive force that drives the potassium-proton antiporters, thus improving active transport to lower internal ionic strength. However, while this adaptation may improve the ability of LD12 cells to live in fresh water, it doesn’t explain why they have lost the ability to survive in salt water where their marine SAR11 cousins thrive.

The most likely explanation comes in the distribution of genes for osmolyte uptake. Critically, LSUCC0530, and LD12 in general, have lost the *proVWX* glycine-betaine/proline ABC transporter shared by all other marine SAR11 (Table 1). Although a relatively minimal number of genes, this uptake capability likely represents a significant loss of function for LD12 salinity tolerance. In *Escherichia coli*, the *proVWX* operon has a strong promoter that upregulates these genes two orders of magnitude in response to increased salinity (Dattananda and Gowrishankar, 1989). Indeed, rapid proline uptake is a common response to osmotic upshift in many bacteria (Empadinhas and da Costa, 2008). Furthermore, LSUCC0530 and other LD12 genomes have lost the ABC transporters for ectoine/hydroxyectoine present in the other SAR11 genomes. Given the slow growth of LD12, rapid production of new intracellular osmolytes in response to increased salinity seems unlikely. Thus, the loss of these transport systems for quickly equilibrating intracellular osmolarity probably prevents successful dispersal of this organism into higher salinity waters.

This finding allows us to speculate further about the evolutionary trajectory that led to successful colonization of freshwater environments by LD12. All evidence suggests that the last common SAR11 ancestor was a marine organism with a streamlined genome (Logares *et al.*, 2010, Luo *et al.*, 2013, Zaremba-Niedzwiedzka *et al.*, 2013). During further diversification into subclade III, additional genes and pathways were lost, for example, ppc and genes for DMSP and some C1 compound metabolisms (Eiler *et al.*, 2016, Sun *et al.*, 2016). Therefore, the common ancestor of subclade IIIa and LD12 would have had to adapt to freshwater environments starting with an already reduced genome and limited metabolic flexibility. A simple way to inhabit lower salinity environments would have been to decrease expression of, or altogether lose, genes for uptake and/or production of osmolytes. If the adaptation were reversible, we would expect to see examples of LD12 organisms growing in marine systems. However, existing evidence suggests that while LD12 organisms can grow (established herein) and be active (Piwosz *et al.*, 2013) in low brackish conditions, they are not successful in higher salinity brackish or marine water (Dupont *et al.*, 2014, Herlemann *et al.*, 2014, Logares *et al.*, 2010, Salcher *et al.*, 2011). Thus, we propose that LD12 adaptation to fresh water involved irreversible loss of function, which decreased osmotic response capability in LD12 cells, thereby preventing recolonization of higher salinity environments. The missing ABC transporters for proline/glycine betaine and ectoine/hydroxyectoine in LD12 are likely candidates for this loss of function. Experimental tests on the differences in production and uptake of osmolytes among subclade IIIa and LD12 representatives will provide critical insight into how these different evolutionary paths have resulted in unique salinity ranges.

For the first cultured representative of the LD12 clade, we propose the provisional taxonomic assignment for strain LSUCC0530 as ‘*Candidatus* Fonsibacter ubiquis’,

### Description of *Fonsibacter* gen. nov

*Fonsibacter* (*Fons* L. noun; fresh water, spring water, - bacter, Gr. adj.; rod, bacterium. Fonsibacter referring to the preferred low salinity habitat of this organism).

Aerobic, chemoorganoheterotrophic, and oligotrophic. Cells are small, curved rods roughly 1 x 0.1μm. Non-motile. Based on phylogenomics and 16S rRNA gene phylogenetics, subclade IIIb/LD12 occurs on a separate branch within the *Pelagibacteraceae* (SAR11), sister to subclade IIIa containing strains HIMB114 and IMCC9063. Due to the depth of branching between these clades using concatenated protein-coding genes, and the 92% and 91% 16S rRNA gene sequence identity with HIMB114 and IMCC9063, respectively, we propose LSUCC0530 and subclade IIIb/LD12 as a novel genus in the *Pelagibacteraceae*.

### Description of *Fonsibacter ubiquis* sp. nov

*Fonsibacter ubiquis* (L. adv. ubiquitous).

In addition to the properties given in the genus description, the species is described as follows. Growth occurs at temperatures between 23°C and 30°C, but not at 11°C or below, nor at 35°C or above. Optimal salinity is 1.45 and below, and growth occurs between 0.36 and 4.68. At optimal temperature and salinity, cells divide at an average rate of 0.52 day^-1^. Genome size is 1.16 Mbp, with 1271 predicted genes, and a GC content of 29.02% (calculated). LSUCC0530 had an ANI of 75.3% and 75.9% and an AAI of 58.1% and 59.7% with HIMB114 and IMCC9063, respectively. The genome is available on GenBank under accession number CP024034.

The type strain, LSUCC0530^T^, was isolated from brackish water from the coastal lagoon Lake Borgne off the coast of southeastern Louisiana, USA.

## Materials and Methods

### Isolation and Identification of LSUCC0530

Isolation, propagation, and identification of strain LSUCC0530 (subclade IIIb) was completed as previously reported (Henson *et al.*, 2016). Briefly, water was collected from the surface of the coastal lagoon Lake Borgne (Shell Beach, Louisiana) (29.872035 N, −89.672819 W) on 1 July, 2016. We recorded salinity on site using a YSI 556 MPS (YSI, Solon, OH, USA) (Table S1). Whole water was filtered through a 2.7 μm filter (Whatman), stained with 1X SYBR Green (Lonza, Basal, Switzerland), and enumerated using the Guava EasyCyte 5HT flow cytometer (Millipore, Massachusetts, USA). After serial dilution, water was inoculated into five 96 x 2.1 mL well PTFE plates (Radleys, Essex, UK) containing 1.7 mL of JW5 medium at an estimated 2 cells well^-1^. Cultures were incubated at room temperature in the dark for three weeks and evaluated for growth using flow cytometry. Positive wells (> 10^4^ cells mL^-1^) were transferred in duplicate to capped, 125 mL polycarbonate flasks (Corning, New York, USA) containing 50 mL of JW5 medium. Upon reaching a density of > 5.0 x 10^5^ cells mL^-1^, cells were filtered onto 25 mm 0.22 μm polycarbonate filters (Millipore, Massachusetts, USA), and DNA extractions were performed using the MoBio PowerWater DNA kit (QIAGEN, Massachusetts, USA) following the manufacturer’s instructions. DNA was amplified as previously reported (Henson *et al.*, 2016) and sequenced at Michigan State University RTSF Genomics Core. Sanger sequences were evaluated using the freely available software 4Peaks (v. 1.7.1)(http://nucleobytes.com/4peaks/). Forward and reverse sequences were assembled using the CAP3 webserver (http://doua.prabi.fr/software/cap3), after reverse reads were converted to their reverse complement at http://www.bioinformatics.org/sms/rev_comp.html. The PCR-generated 16S rRNA gene sequence for strain LSUCC0530 is accessible on NCBI GenBank under the accession number KY290650.

### Community iTag analysis

Community DNA from 17 sites of various salinities was filtered, extracted, and analyzed following our previously reported protocol (Henson *et al.*, 2016). The sites sampled were Lake Borgne (LKB; Shell Beach, LA), Bay Pomme D’or (JLB; Buras, LA), Terrebonne Bay (TBon; Cocodrie, LA), Atchafalaya River Delta (ARD; Franklin, LA), Freshwater City (FWC; Kaplan, LA), Calcasieu Jetties (CJ; Cameron, LA). Each site was visited three times with the exception of TBon, which was visited only twice. Briefly, extracted DNA was sequenced at the 16S rRNA gene V4 region (515F, 806R) (Parada *et al.*, 2015), with Illumina MiSeq 2x250bp paired-end sequencing at Argonne National Laboratories. 16S rRNA gene amplicon data was analyzed and classified (OTU0.03) with Mothur v.1.33.3 (Schloss *et al.*, 2009) using the Silva v119 database (Pruesse *et al.*, 2007). To determine the best matching OTU for LSUCC0530, sequences from the OTU representative fasta files, provided by mothur using *get.oturep*(), were used to create a blast database (*formatdb*) against which the LSUCC isolate 16S rRNA genes could be searched via blastn (BLAST v 2.2.26). All best hits ranked at 100% identity. OTU abundance analysis was conducted within the R statistical environment v3.2.3, within the package PhyloSeq (v1.14.0) (McMurdie and Holmes, 2013). Sequences were rarefied using the command *rarefy_even_depth*() and then trimmed for OTUs with at least 2 reads in at least 90% of the samples. Abundances of an OTU were averaged across biological duplicates and a rank abundance matrix was calculated using the command *transform_sample_counts*() with the argument function of function(x) x / sum(x). Completed rarefied rank abundance OTU tables were plotted using the graphing program GGPLOT2 (v. 2.0.0) (Wickham, 2011). All iTag sequences are available at the Short Read Archive with accession number SRR6235382-SRR6235415.

### Growth experiments

Salinity tolerance was tested by growing LSUCC0530 in different media distinguished by proportional changes in the concentration of major ions. All other media elements (carbon, iron, phosphate, nitrogen, vitamins, and trace metals) were kept consistent. Media included JW1, JW2, JW3, and JW4, previously published (Henson *et al.*, 2016), and JW3.5, JW4.5, JW5, and JW6. Recipes for all media used in this study are provided in Table S1. Salinity tolerance experiments were conducted in quintuplicate at room temperature. Temperature range and sole carbon substrate use were tested using the isolation medium, JW5. Temperature range experiments were also conducted in quintuplicate. For all experiments, growth was measured using flow cytometry as described above.

### Microscopy

Strain LSUCC0530 cells were grown to near max density (1x10^7^ cells mL^-1^) in JW5 medium. Cells for transmission electron microscopy were prepared as previously described (Rappé *et al.*, 2002) with one minor change: centrifuged cells were concentrated in JW5 medium with no added fixative. Cells were visualized under a JEOL JSM-2011 High Resolution transmission electron microscope at the LSU Socolofsky Microscopy Center, Baton Rouge, LA. For scanning electron microscopy, cultured cells were fixed for four hours in 4% Glutaraldehyde buffered with 0.2M Cacodylate buffer and filtered onto a 0.2 μm nylon filter (Pall, Michigan, USA). Samples were dehydrated in an ethanol series (30%, 50%, 70%, 96 and 100%) for 15 minutes each, followed by critical point drying. Cells were visualized under a JEOL JSM-6610 scanning electron microscope at the LSU Socolofsky Microscopy Center, Baton Rouge, LA.

### Genome sequencing, assembly, and annotation

Cells were grown in 1 L JW5 medium (Table S1) and filtered onto 0.2 μm nylon filters (Pall, Michigan, USA). DNA was obtained via phenol-chloroform extraction and sequenced using both Illumina HiSeq and MiSeq approaches. *HiSeq*. 350 ng of DNA was sheared to ~500 bp with a Qsonica Q800R (QSonica) using 16 cycles with a amplitude of 25% and pulse rate of 20 seconds on, 20 seconds off. DNA was visualized on a 1.5% agrose gel stained with Gel Red (Biotum) to determine average sheared length and quantified using Qubit high sensitivity dsDNA assay kit (Invitrogen). Library prep was conducted following a modified Kapa Hyper Prep Kit protocol (Kapa Biosystems, Inc.). Briefly, we ligated end-repaired and adenylated DNA to universal Y-yoke oligonucleotide adapters and custom Illumina iTru dual-indexed primers (Glenn *et al.*, 2016). Prior to index amplification, post-ligation LSUCC0530 DNA was cleaned using 1.2x SPRI (Roland and Reich, 2012). Following a brief, 8 cycle PCR amplification of the library to add indexes, the genomic library was quantified using a Qubit broad range dsDNA assay kit (Invitrogen). Illumina HiSeq 3000 sequencing was performed at Oklahoma Medical Research Facility using SBS kit chemistry to generate 150 bp paired-end reads, insert size 400 bp. *MiSeq*. DNA was sent to the Argonne National Laboratory Environmental Sample Preparation and Sequencing Facility which performed library prep and sequencing, generating 250 bp paired-end reads, insert size 550 bp.

Due to the large number of reads from the HiSeq sequencing, we created a random subset using 14% of the total reads by using the program *seqtk* (v. 1.0-r75-dirty) and the flag –s100. We trimmed reads for adapter contamination and quality using Trimmomatic (Bolger *et al.*). Initial assembly with the subset of reads was done with SPAdes (Bankevich *et al.*, 2012). This generated a large single scaffold with overlapping ends. We performed a quality assessment of the single scaffold via iterations of Reapr (Hunt *et al.*, 2013)-advised scaffold breaks and SSPACE (Boetzer *et al.*, 2011) extensions using both the full set of HiSeq reads and the MiSeq reads. Final assessment of the scaffold with overlaps removed from the ends was performed with Pilon (Walker *et al.*, 2014) using all *HiSeq* reads mapped to the scaffold with BWA (Li and Durbin, 2009). No issues were reported. An analysis of the GC skew was performed on the final scaffold using the gc_skew script provided via Brown et al. 2015 (Brown *et al.*, 2015). Contamination and completion were also evaluated with CheckM (Parks *et al.*, 2015). Detailed commands and outputs for the assembly and quality assessment are provided in Supplementary Information. The final circular chromosome scaffold was annotated by IMG (Markowitz *et al.*, 2014) and is publicly available with IMG Taxon ID number 2728369501. It is also publicly available in GenBank with accession number CP024034. Genome sequencing reads are available in SRA with accession numbers SRR6257553 (MiSeq) and SRR6257608 (HiSeq).

### Comparative genomics

Orthologus clusters were determined using the Get Homologues pipeline (Contreras-Moreira and Vinuesa, 2013) using the LSUCC0530 genome and 48 publicly available SAR11 genomes from IMG, including the 10 LD12 SAGs analyzed previously (Eiler *et al.*, 2016, Zaremba-Niedzwiedzka *et al.*, 2013) (Table S1). Clusters were determined from amino acid sequences using the OrthoMCL option in Get Homologues. All clusters are provided in cluster_list.txt as part of Supplemental Information. Average amino acid identity (AAI) and synteny were calculated as previously reported (Dick *et al.*, 2009). Average nucleotide identity (ANI) was calculated with the Konstantinidis Lab website ANI calculator (http://enve-omics.ce.gatech.edu/ani/index) as reported (Goris *et al.*, 2007). Compatible solutes identified in Table S1 were identified in SAR11 genomes using KO numbers, annotations, and cross-referenced with orthologous clusters.

### Bacteriophage searches

We looked for signatures of bacteriophage in the LSUCC0530 genome using two methods, VirSorter (Roux *et al.*, 2015) and PHASTER (Arndt *et al.*, 2016) with default settings. CRISPR regions are identified as part of the standard IMG annotation process, and none were found in LSUCC0530.

### Phylogenomics

We selected single-copy genes present in at least 39 of the 49 genomes as determined via Get Homologues (above). Amino acid sequences from these 83 markers were separately aligned with MUSCLE (Edgar, 2004), culled with Gblocks (Castresana, 2000), and concatenated into a superalignment totaling 23,356 positions. Inference was performed with RAxML (Stamatakis *et al.*, 2008) using the PROTCATLG model and 100 bootstrapping runs. All scripts (with settings) used in the alignment, culling, concatenation, and inference processes are available in Supplemental Information. Node labels were renamed with Newick Utilities (Junier and Zdobnov, 2010) *nw_rename*, and the tree was visualized with Archaeopteryx (Han and Zmasek, 2009).

### Single gene phylogeny

The 16S rRNA gene sequence from the LSUCC0530 genome was aligned with 284 alphaproteobacterial sequences and seven outgroup sequences. We completed phylogenetic inference for *aceA* and *malA* in an analogous manner. For each, the LSUCC0530 gene was searched against the NCBI nr database using blastp (v. 2.2.28). The top 100 sequence hits were combined with all homologus SAR11 sequences identified with Get Homologues for the *aceA* phylogeny, and both the *malA* and *malG* sequences in all SAR11s for the *malA* phylogeny. Sequences from redundant taxa were removed. For all trees, alignment with MUSCLE (Edgar, 2004), culling with Gblocks (Castresana, 2000), and tree inference with FastTree2 (Price *et al.*, 2010) was completed with the FT_pipe script, provided in Supplemental Information, along with the initial fasta file containing all sequences and the newick tree output file. Node labels were changed with Newick Utilities (Junier and Zdobnov, 2010) *nw_rename* and visualization was performed with Archaeopteryx (Han and Zmasek, 2009).

### Metagenomic recruitment

Scaffolds for the 49 SAR11 genomes, including strain LSUCC0530, used in the phylogenomic analyses were used for competitive recruitment of metagenomic sequences from 89 different samples. Prior to recruitment, multi-scaffold assemblies were condensed into single scaffolds, and all single scaffolds were used to generate a BLAST database with the command *makeblastdb* (BLAST v. 2.2.28). Metagenomic datasets from 90 different environmental samples of various salinities from the Baltic Sea (Dupont *et al.*, 2014), Amazon River, freshwater lakes (Eiler *et al.*, 2014, Martinez-Garcia *et al.*, 2011), Lagoon Albufera (Ghai *et al.*, 2012), Columbia River (Fortunato and Crump, 2015, Smith *et al.*, 2013b), Lake Lanier (Oh *et al.*, 2011), Lake Michigan (Denef *et al.*, 2016), Feitsui Reservoir (Tseng *et al.*), and GOS (Rusch *et al.*, 2007) were downloaded from the Short Read Archive (SRA) and European Nucleotide Archive (ENA) using *ftp* and SRA toolkit (v.2.8.2-1), respectively. Reads from pyrosequencing were dereplicated with the program CD-HIT (v 4.6) (Fu *et al.*, 2012) using the *cd-hit-454* command and flag - c 0.95. Illumina sequence data was converted from fastq to fasta files with the FastX toolkit (v. 0.0.13.2) (http://hannonlab.cshl.edu/fastx_toolkit) using the command *fastq_to_fa*sta with the flag - Q33. Metagenomic sequences were separated into 1 million sequence files and recruited against the SAR11 database with blastn (v. 2.2.28) with –max_target_seqs 1. Best hits for each metagenomic sequence were kept if the alignment length was > 90% of the median length of all metagenomic sequences for a given sample. Metagenomic recruitment was expressed in read density per genome using ggplot2 *geom_density*() and reads per kilobase of genome per million mapped reads (RPKM) as previously reported (Thrash *et al.*, 2017).

### Script availability

All scripts used in this work can be found on the Thrash Lab website with the manuscript link at http://thethrashlab.com/publications.

## Acknowledgements

Funding for this work was provided in part by a Louisiana Board of Regents (grant LEQSF(2014-17)-RD-A-06) grant to JCT and the LSU Department of Biological Sciences (startup funds to JCT and BCF). The authors thank the LSU Socolofsky Microscopy Center for assistance with electron microscopy imaging. Portions of this research were conducted with high performance computing resources provided by Louisiana State University (http://www.hpc.lsu.edu). The authors also thank Katherine D. McMahon and Sarah L. R. Stevens for helpful commentary on the manuscript draft.

## Author Contributions

JCT designed the study; MWH conducted cultivation experiments, nucleic acid extraction, microscopy, and iTag and metagenomics analyses; JCT and BCF assembled the genome and conducted quality assessment; MWH, VCL, and JCT conducted comparative genomics analyses, metabolic reconstruction, phylogenetic, and phylogenomic analyses; JCT lead manuscript writing and all authors contributed to and reviewed the text.

## Conflict of Interest

The authors declare no conflict of interest.

